# Contrasting effects of commercial and native arbuscular mycorrhizal fungal inoculants on plant biomass allocation, nutrients and phenolics

**DOI:** 10.1101/2020.04.28.065748

**Authors:** Adam Frew

## Abstract

As the global population increases, the need to feed more people must be met while simultaneously conserving the long-term sustainability of our agroecosystems. There is mounting interest and discussion around the application of arbuscular mycorrhizal fungal (AMF) inoculants to enhance crop growth, nutrient uptake and pest resistance. However, the effects of AMF inoculation are variable and context dependent. This study shows the stronger effects of an AMF inoculant with greater number of fungal species, but that these effects are no better than re-inoculating plants with a field-sourced native AMF inoculant.

## Introduction

Global food security relies heavily on a select number of plant species, most of which associate with arbuscular mycorrhizal fungi (AMF) of the subphylum, Glomeromycotina (Smith & Smith, 2011). Arbuscular mycorrhizas provide plants with an array of functions including nutrient acquisition and protection from abiotic and biotic stresses. These fungi also play an important role in many ecosystem level processes, contribute to soil structure and health, and have strong effects on plant community ecology (Tedersoo *et al.*, 2020).

Given the capacity for the AM symbiosis to provide ecological and crop benefits, and the serious concern of global soil ‘health’, there is increasing recognition of the importance of managing AMF to the future of food production (Thirkell *et al.*, 2017; Rillig *et al.*, 2019). One aspect of this is the application of AMF inocula to encourage mycorrhization of crops. However, the outcome of engaging in the AM symbiosis can be highly context dependent, subject to AMF and plant species identities, and on local soil conditions. For example, nutrient exchange between fungus and plant can vary between crop cultivar (Elliott *et al.*, 2020), and studies show a certain level of partner selectivity exists in these plant-fungal associations (Sepp *et al.*, 2019). Additionally, evidence suggests that certain AMF taxa may be more associated with particular functions such as plant nutrient uptake, or plant resistance against pests and pathogens (Bennett & Bever, 2007; Wehner *et al.*, 2010). Indeed, AMF have been shown to differentially affect plant secondary metabolites associated with resistance to insect herbivores including phenolics (Mithöfer & Boland, 2012) and benzoxazinoids (Frew *et al.*, 2018).

Despite evidence of context-dependent functional diversity accross AMF taxa, there are relatively few examinations at the fungal community level. Indeed, different combinations of AMF taxa differentially interact and can exhibit functional complementarity (Jansa *et al.*, 2008; Sikes *et al.*, 2010). For example, studies have shown that inoculants containing more than one AM fungal species can have stronger or weaker effects compared to single species inoculants (Veresoglou *et al.*, 2012; Grümberg *et al.*, 2015). Yet, our understanding of how assemblages of AMF communities (including species richness) might correlate with different crop nutritional and stress resistance traits remains ambiguous at best. Consequently, it is a gamble whether the AMF taxa in a given inoculum will provide the wanted outcomes, or are indeed ‘superior’ to the native fungal community already present in the soil (Hart *et al.*, 2018).

Therefore this study examined the effects of inoculation with a single AM fungal species, a combination of four AM fungal species, and a native field soil inoculum. The effects on plant biomass allocation, nutrient uptake (phosphorus and nitrogen), and a group of resistance-associated metabolites (phenolics) were assessed in two globally significant crop species.

## Methods

### Experimental set-up

*Hordeum vulgare* L. cv. ‘Hindmarsh’ (90 plants) and *Sorghum bicolor* L. Moench cv. ‘Enforcer’ (90 plants) were grown in 3.7L pots, one plant per pot, with gamma-irradiated 80: 20 soil: quartz sand mixture (Table S1; see Supporting Information for more detailed methodology). Plants were grown under one of three AMF treatments (by directly pipetting ~400 spores onto roots) which comprised of either (i) one AMF species from a commercial inoculum containing *Rhizophagus irregularis*; (ii) four AMF species from a commercial inoculum containing *Claroideoglomus etunicatum*, *Funneliformis coronatum*, *F. mosseae* and *Rhizophagus irregularis*; (iii) native AMF community comprising AMF spores extracted from the field soil. All pots received microbial filtrate (300 ml) to standardise the microbial community across pots. Plants were grown in a growth chamber (Conviron® PGW40) with day: night air temperatures of 27 °C and 17 °C (±4 °C) respectively, daylight set at 900 mol^−2^s^−1^ on a 12h photoperiod. Every two weeks pots were rearranged within the chamber to reduce any spatial effects.

After ten weeks plant roots were washed and a 1-2g subsample of fine roots were taken from a random selection of 10 plants per treatment from each plant species for mycorrhizal colonisation scoring. Aboveground tissue was snap frozen in liquid nitrogen before being freeze dried prior to chemical analysis.

### Plant chemistry and fungal colonisation

Root subsamples were cleared with 10% KOH and stained with 5% ink-vinegar (Vierheilig *et al.*, 1998). Mycorrhizal colonisation was assessed using the gridline-intersect method with at least 100 intersects per sample (McGonigle *et al.*, 1990). Freeze-dried and ground plant material was analysed for nitrogen concentrations using an elemental analyser (LECO TruMac CNS analyser, LECO, Saint Joseph, MI, USA) and for phosphorus concentrations using inductively coupled plasma-optical emission spectrometer (ICP-OES) (Varian 710-ES; Agilent Technologies Inc., Palo Alto, CA, USA).

### Statistics

R statistical interface (v3.6.1) was used for all statistical analysis.

Data exploration for all responses was carried out following the protocol described in Zuur, Ieno, and Elphick (2010). The effects of the AMF treatments on measured parameters of the two plant species were assessed by fitting standard linear models using the *lm* function comparing factors ‘species’, ‘AMF’ and their interactions, then applying *Anova* function from the R package ‘car’ (Fox & Weisberg, 2011). To satisfy model assumptions, belowground biomass, aboveground biomass and N:P were transformed to give residual diagnostic plots which fit a normal distribution. Tukey post-hoc tests using the *HSD.test* function from the R package ‘agricolae’ (De Mendiburu, 2019) were used to identify statistical differences between groups. Nitrogen concentration and vesicular colonisation response variables were analysed by fitting generalised linear models (family = poisson) using the *glm* function followed by chi-squared test using the *Anova* function from the R package ‘car’ (Fox & Weisberg, 2011).

## Results

*Hordeum* sp. plants inoculated with four AMF and the native AMF had 25% and 22% lower root:shoot, respectively, compared with those inoculated with a single AMF species (Table S1, Figure 1a). This was largely driven by reductions in belowground biomass (Figure S1b).

**Figure 1.**
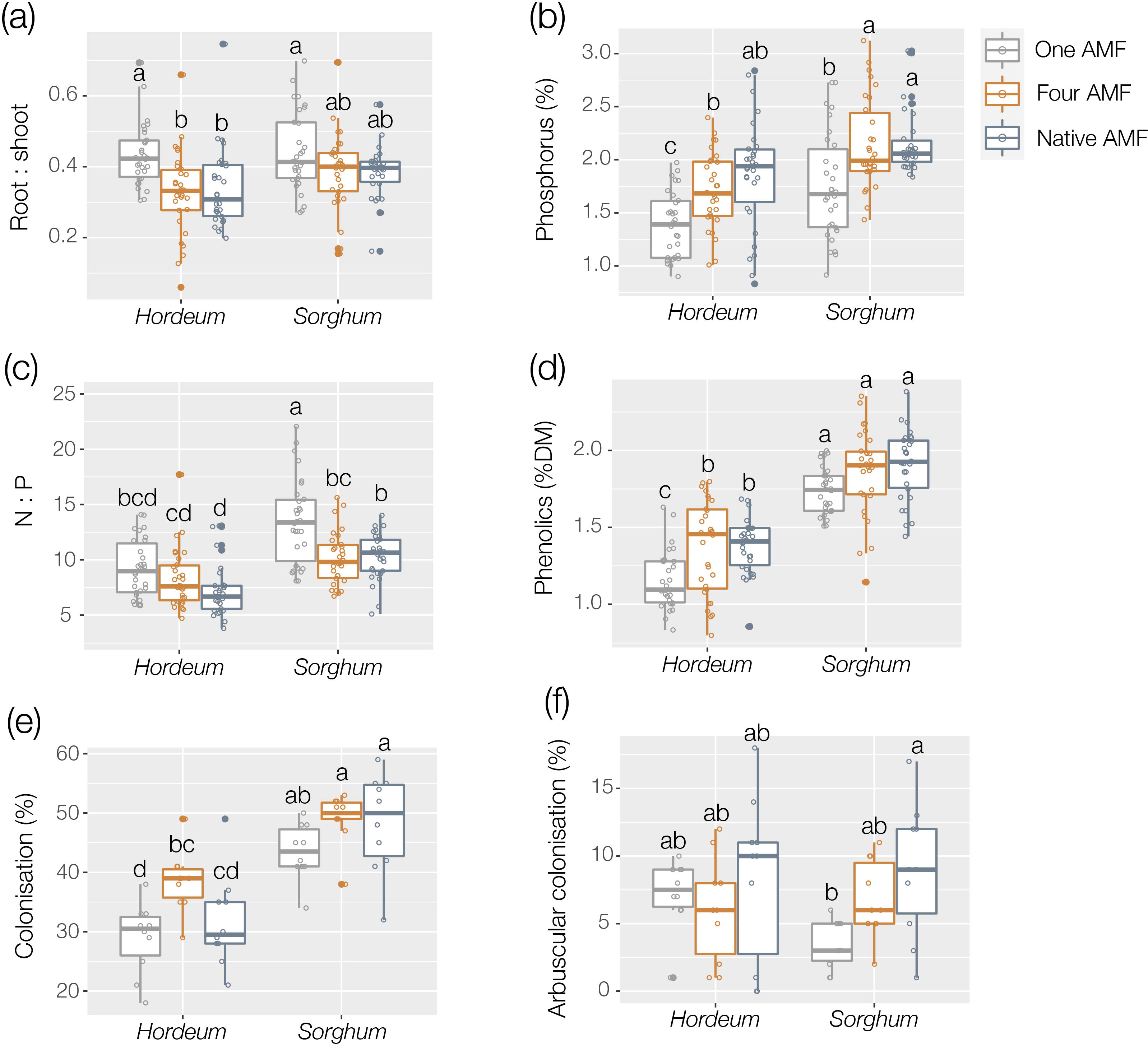
Effects of inoculation with one arbuscular mycorrhizal fungal (AMF) species, four AMF species or with a native AMF inoculant extracted from field soil on the **(a)** root:shoot, **(b)** phosphorus concentration (%), **(c)** N:P, **(d)** total phenolics (%DM), **(e)** total AMF root colonisation (%), and **(f)** arbuscular root colonisation (%) in *Hordem vulgare* L. cv. ‘Hindmarsh’ and *Sorghum bicolor* L. Moench cv. ‘Enforcer’. Different letters indicate boxes that are significantly different from each other (*P*<0.05, Tukey HSD).

*Hordeum* sp. phosphorus concentrations were 24% and 35% greater in plants inoculated with four AMF and the native AMF, respectively, compared to those inoculated with one AMF (Figure 1b). *Sorghum* sp. displayed a similar response with 22% and 23% greater phosphorus concentrations in plants inoculated with four AMF and native AMF, respectively, compared to the one AMF inoculant (Figure 1b). These effects on phosphorus concentrations in *Sorghum* sp. were somewhat reflected in foliar N:P which was significantly lower in plants treated with four AMF and native AMF inocula compared with those plants under the one AMF treatment (Figure 1c). In contrast, the AMF inocula did not differentially affect N:P in *Hordeum* sp. plants. Foliar nitrogen differed overall between the two plant species but was unaffected by the AMF treatments (Table S1, Figure S1c).

*Sorghum* sp. had 41% more foliar phenolics than *Hordeum* sp. (Table S1, Figure 1d). Phenolic concentrations did not differ between AMF treatments in Sorghum spp., while *Hordeum* sp. plants inoculated with four AMF species and the native AMF had higher phenolic concentrations than plants inoculated with one AMF (Table S1, Figure 1d)

Overall, total AM fungal root colonisation was 42% higher in *Sorghum* sp. compared with *Hordeum* sp. (Table S1, Figure 1e). Total colonisation only differed between the different AMF inocula in *Hordeum* sp. roots (Figure 1e), while formation of arbuscules differed between AMF inocula in *Sorghum* sp. roots (Figure 1f).

## Discussion

This study found that inoculation with four AMF species had stronger effects on plant allometric partitioning, foliar nutrient and phenolic concentrations than inoculation with a single AMF species. This finding is generally consistent with previous studies where inoculants with greater AMF species richness tend to have stronger effects on different host plant traits of interest (Jansa *et al.*, 2008; Veresoglou *et al.*, 2012; Frew, 2019). However, the results here also show that the effects of inoculating with four AMF species were no different from the effects of applying a native AMF inoculant, extracted from field soil. Thus, the application of commercial AMF inocula to soil may not deliver effects that are not already obtainable from the resident AMF community present in the environment. Yet, these results also point out that AMF incoula may provide significant benefits to plants grown in substrates with impoverished AMF diversity.

Although inoculating with four AMF species or the native AMF had similar outcomes, the effects differed between the two crop species. For example, in *Hordeum* sp. the four and native AMF treatments reduced root:shoot, and increased phosphorus and phenolics compared to the inoculant with a single AMF species. Contrastingly, in *Sorghum* sp. the four and native AMF treatments did not affect root:shoot ratio between AMF treatments, but did increase phosphorus and reduce foliar N:P compared to the one AMF species treatment.

The biomass allocation away from the roots observed here is a commonly reported effect of the AM symbiosis (Veresoglou *et al.*, 2012), which can be attributed to improved nutrition. Although the root to shoot ratio is a relatively crude measure, it is proposed that biomass investment towards roots decreases as nutrient requirements are met. Although root:shoot did not differ between AMF treatments in *Sorghum* sp., both plant species exhibited greater phosphorus concentration under the four and native AMF treatments compared to inoculation with one AMF. Thus, it is notable that N:P was reduced by the four and native AMF treatments in *Sorghum* sp. and not *Hordeum* sp. as the reduced biomass allocation towards the roots observed in *Hordeum* sp. under these same treatments might have otherwise suggested nutrient limitation under the one AMF treatment.

The increased phenolics in *Hordeum* sp. under the four and native AMF compared to the one AMF treatment is also noteworthy. Previous studies report increases in phenolics from the AM symbiosis (Jung *et al.*, 2012), yet this is the first evidence, to my knowledge, that inoculation with different AMF communities differentially affects phenolics between plant species. Although a relatively simplistic measure, total phenolics are associated with resistance to insect herbivory (Mithöfer & Boland, 2012). Thus, the findings here call for a more detailed examination of how differences in AMF community assembly affect phenolic-based resistance to herbivory.

Despite controversies around AMF inoculants and the variability of their efficacy, the management of mycorrhizal fungi is likely to have an increasingly important role in future sustainable food production. Although this study was under controlled conditions, the results presented here highlight that the application of multispecies AMF inocula can have beneficial outcomes for the host plants, but also that inoculant AMF communities may provide little to no additional benefit compared with the resident AMF community. Our knowledge around effectively managing the AM symbiosis in plant production systems is still developing and therefore practitioners should take a thoughtful and well-considered approach when it comes to applying AMF inoculants in the field.

## Acknowledgements

The author would like to thank the technical teams at Charles Sturt University for their support including the technical team in the National Life Sciences Hub, in particular Vincent West and Yameng Tian. Funding and support was provided by a Charles Sturt University Faculty of Science Postdoctoral Research Fellowship awarded to AF.

