## Supporting Information for "Contrasting effects of commercial and native arbuscular mycorrhizal fungal inoculants on plant biomass allocation, nutrients and phenolics"

**Author:** Adam Frew

The following Supporting Information is available for this article:

**Extended experimental set-up methods** Extended methods outlining experimental set-up, plant growth, harvest and sample collection.

**Table S1** Nutrient analysis of fully homogenised soil substrate.

**Table S2** Results of ANOVAs on fitted standard linear models, and chi-squared tests on fitted generalised linear models, both examining main effects and interactions of factors ‘Species’ and ‘AMF’ on plant biomass and chemical response variables. Significant impacts (P < 0.05) are indicated in bold.

**Figure S1** Effects of inoculation from one AMF, four AMF or a native AM fungal community on the biomass (g), nitrogen concentration (mg g^-1^) and vesicular colonisation (%) in Hordeum vulgare L. cv. ‘Hindmarsh’ and Sorghum bicolor L. Moench cv. ‘Enforcer’.

**Extended experimental set-up methods**

*Hordeum vulgare* L. cv. ‘Hindmarsh’ (90 plants) and *Sorghum bicolor* L. Moench cv. ‘Enforcer’ (90 plants) were grown in 3.7L pots, one plant per pot, with 3.42 kg (oven dry equivalent) of gamma irradiated 80 : 20 soil : quartz sand (glass grade sand, Australian Silica Quartz Pty. Ltd.) mixture (see Supporting information and Table S3 for soil details growth media description). Soil was collected from previously grazed paddock site (142 m a.s.l. elevation) in the Riverina region of NSW, Australia (35°02'45.6"S 147°20'53.8"E). Thus, 30 plants of each species were grown under one of three AM fungal treatments of approx. 400 fungal spores by pipetting directly onto seedling roots. All spores were extracted from the inoculum using the wet sieving and sucrose centrifugation method (Daniels & Skipper, 1982). The treatments comprised of either (i) **one AMF** species using a commercial inoculum (Microbe Smart Pty. Ltd. Melrose Park DC. South Australia) containing only *Rhizophagus irregularis*; (ii) **four AMF** species using a commercial inoculum (Microbe Smart Pty. Ltd. Melrose Park DC. South Australia) containing four species identified as *Claroideoglomus etunicatum*, *Funneliformis coronatum*, *F. mosseae and* *Rhizophagus irregularis*; (iii) **native AMF** community comprising AM fungal spores extracted from the field soil. All pots received microbial filtrate (300 ml) made of equal parts of extraneous extraction solution (i.e. without AM fungal spores) from the three treatments to standardise the background microbial community within each pot. Plants were grown in a growth chamber (Conviron® PGW40) with day : night air temperatures of 27 °C and 17 °C (±4 °C) respectively, daylight set at 900 mol^-2^s^-1^ on a 12h photoperiod and relative humidity controlled at 60% (±8%). Every two weeks pots were rearranged within the chamber to reduce any spatial effects.

After ten weeks all plants were carefully removed from their pots, their roots were washed and a 1-2g subsample of fresh fine roots were taken for mycorrhizal colonisation scoring, while aboveground plant tissue was removed and snap frozen in liquid nitrogen before being freeze dried for 42 hours prior to chemical analysis.

**Table S1**. Analysis of soil collected from previously grazed paddock site (142 m a.s.l. elevation) in the Riverina region of NSW, Australia (35°02'45.6"S 147°20'53.8"E). Nutrient analysis of fully homogenised soil substrate. Analysis carried out by Environmental Analysis Laboratory, Southern Cross University, Lismore, Australia.

| **Nutrient (Method)** | **Units** | **Soil** |
| --- | --- | --- |
| pH (water) | pH unit | 5.14 ± 0.30 |
| Exchangeable magnesium (ammonium acetate) | mg/kg | 179.5 ± 1.71 |
| Exchangeable calcium (ammonium acetate) | mg/kg | 785.75 ± 13.97 |
| Exchangeable potassium (ammonium acetate) | mg/kg | 304.25 ± 13.29 |
| Exchangeable sodium (ammonium acetate) | mg/kg | 26.5 ± 1.5 |
| Ammonium (KCl) | mg/kg | 12.28 ± 2.43 |
| Total nitrogen (LECO analyser) | % | 0.05 ± 0.01 |
| Total carbon (LECO analyser) | % | 0.56 ± 0.12 |
| Total phosphorus (acid extractable) | mg/kg | 254.75 ± 5.07 |
| Phosphorus (colwell) | mg/kg P | 32 ± 3.34 |
| Phosphorus (Bray1) | mg/kg P | 13.03 ± 2.92 |

**Table S2** Results of ANOVAs on fitted standard linear models and chi-squared tests on fitted generalised linear models examining main effects and interactions of factors ‘Species’ and ‘AMF’ on plant biomass, plant chemistry, and AMF colonisation response variables. Significant impacts (P < 0.05) are indicated in bold.

|  | Species | | AMF | | Species x AMF | |
| --- | --- | --- | --- | --- | --- | --- |
|  | F _1,174_ | *P* | F _2, 174_ | *P* | F _2, 174_ | *P* |
| Total biomass | **298.09** | **<0.001** | 0.246 | 0.782 | 2.019 | 0.136 |
| Aboveground biomass † | **222.74** | **<0.001** | 0.600 | 0.549 | 2.226 | 0.111 |
| Belowground biomass † | **242.16** | **<0.001** | **8.635** | **<0.001** | 0.716 | 0.49 |
| Root : shoot | **6.588** | **0.011** | **11.52** | **<0.001** | 1.017 | 0.364 |
| Phosphorus | **35.65** | **<0.001** | **20.09** | **<0.001** | 0.485 | 0.617 |
| Nitrogen ‡ | **195.0** | **<0.001** | 0.633 | 0.729 | 3.36 | 0.186 |
| N : P § | **51.159** | **<0.001** | **10.465** | **<0.001** | 1.313 | 0.272 |
| Phenolics | **249.51** | **<0.001** | **13.039** | **<0.001** | 0.62 | 0.539 |
|  | F _1,54_ | *P* | F _2, 54_ | *P* | F _2,54_ | *P* |
| Total colonisation | **75.13** | **<0.001** | **7.45** | **0.002** | 1.20 | 0.308 |
| Arbuscular colonisation | 0.609 | 0.438 | **3.585** | **0.035** | 2.146 | 0.127 |
| Vesicular colonisation ‡ | 0.014 | 0.906 | 2.808 | 0.246 | 2.762 | 0.251 |

† Sqrt transformed; § log transformed; ‡ Generalized linear model (family = poisson)

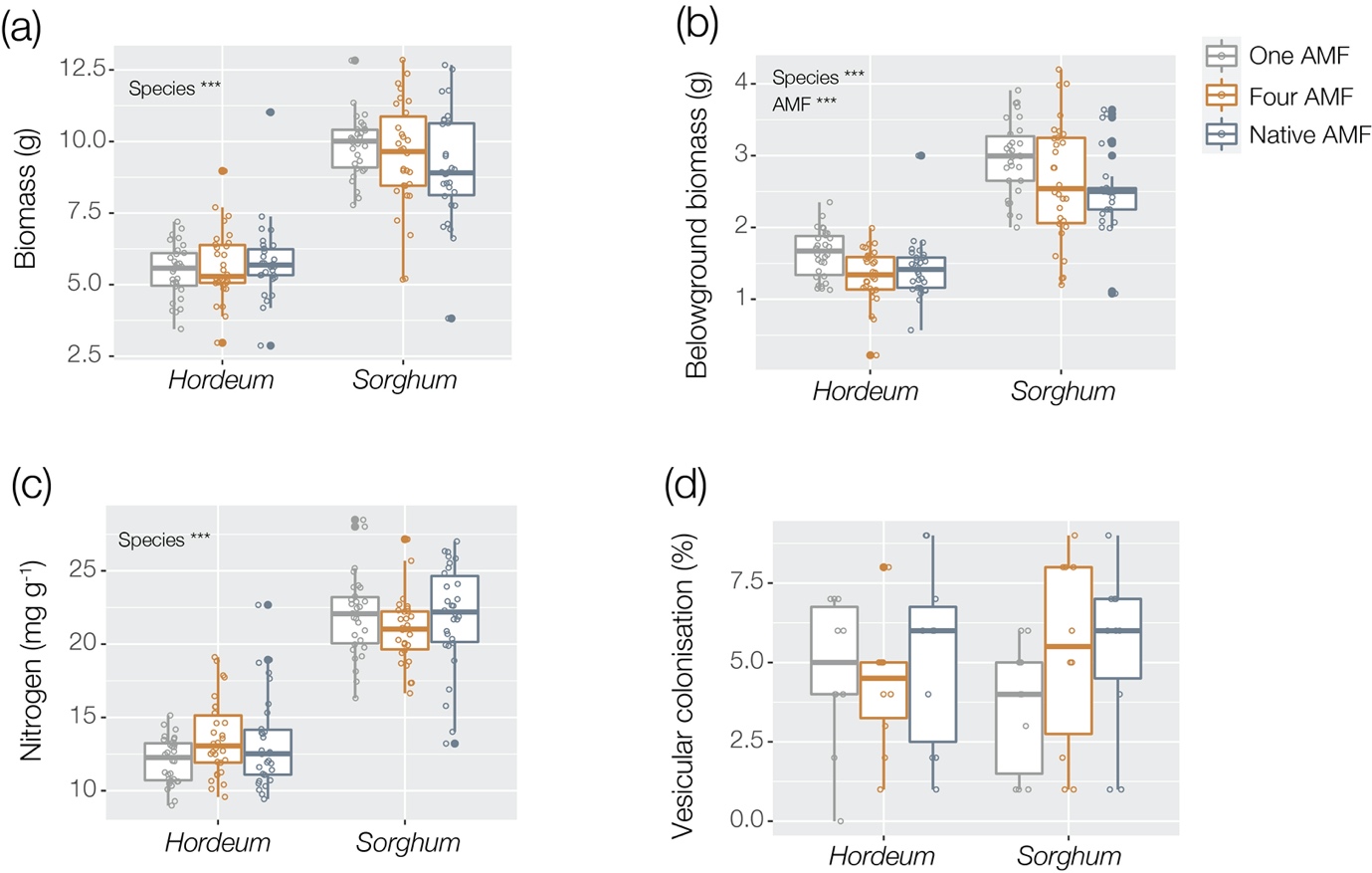

**Figure S1** Effects of inoculation from one AMF, four AMF or a native AM fungal community on the **(a)** biomass (g), **(b)** belowground biomass (g), **(c)** nitrogen concentration (mg g^-1^) and **(d)** vesicular colonisation (%) in *Hordeum vulgare* L. cv. ‘Hindmarsh’ and *Sorghum bicolor* L. Moench cv. ‘Enforcer’. Significant factors are shown, degrees of significance are indicated as follows: *** *P* < 0.001
